# Delayed Anxiolytic-Like Effects of Psilocybin in Male Mice Are Supported by Acute Glucocorticoid Release

**DOI:** 10.1101/2020.08.12.248229

**Authors:** Nathan T Jones, Zarmeen Zahid, Sean M Grady, Ziyad W Sultan, Zhen Zheng, Matthew I Banks, Cody J Wenthur

**Author notes:** **Corresponding Author** Cody J Wenthur, Mailing Address: Rennebohm Hall, 777 Highland Ave, Madison, WI 53705.

## Abstract

Despite observed correlations between acute glucocorticoid release, self-reported anxiety, and long-term treatment outcomes for human studies using psilocybin-assisted psychotherapy approaches, the mechanistic relationship between psychedelic-dependent stress and subsequent behavioral responses remains unclear. Using rodents, direct manipulation of stress-associated hormone responses can be achieved with established pharmacologic models for the assessment of antidepressant and anxiolytic therapeutics. Here, chronic oral corticosterone-induced suppression of the hypothalamic-pituitary-adrenal axis is used to assess the relevance of drug-induced glucocorticoid release on the acute, post-acute, and long-term effects of psilocybin in male C57BL/6J mice. In these studies, psilocybin-induced acute anxiogenesis was found to be correlated to post-acute anxiolysis in a dose-dependent manner. Psilocybin also displayed acute increases in plasma corticosterone, but a post-acute anxiolytic effect in the novelty suppressed feeding test. Both effects were lost when psilocybin was administered in animals pre-exposed to chronic oral corticosterone. A similar long-term interaction between chronic corticosterone and psilocybin administration was observed in an open field test occurring one week after drug administration. Psilocybin administration alone led to more time spent in the center of the arena, but animals spent less time in the center with chronic corticosterone exposure. Intriguingly, these interactive effects were absent in animals exposed to brief isoflurane anesthesia after drug treatment. Overall, these experiments identify acute glucocorticoid release as a relevant biological modifier for the post-acute and long-term behavioral effects of psilocybin in mice. Rodent studies are thus suggested as a tractable means to address neuroendocrine mechanisms supporting context-dependent psychedelic effects in mammalian species.

## 1. Introduction

Investigations applying the serotonin 2A receptor (5-HT_2A_R) agonist psilocybin as a psychiatric medication have recently proliferated.[1–4] This classical psychedelic compound has been investigated in multiple clinical trials for substance use disorders (SUD), end-of-life anxiety, and major depressive disorder (MDD), including treatment resistant forms.[5–10] However, all modern clinical studies of therapeutic interventions with psychedelics are intrinsically measuring the overall effects of an intervention package that includes intensive psychological preparation, debriefing sessions, as well as guided support and environmental manipulations during the administration session.[11,12].

Animal models may provide a means to help isolate the pharmacologically-mediated effects of psychedelics in the absence of additional psychological support. One important example of such direct biological effects is the demonstrated ability of classical psychedelics to induce both functional and structural plasticity in neuronal preparations from mammalian and non-mammalian sources.[13–19] Additional studies assessing antidepressant-like and anxiolytic-like effects following treatment with 5-HT_2A_R agonists in rodent behavioral tests like the Forced Swim Test (FST), Open Field Test (OFT), and Elevated Plus Maze (EPM) have also emerged more recently. [19–26] The observed results have sometimes been consistent with interpretations of antidepressant-like or anxiolytic-like activity, such as reductions in FST immobility time and increased open arm time in the EPM. However, there have also been results that are inconsistent with this interpretation, such as a lack of effect on FST in Flinders’ Sensitive Rats for up to a week after single or repeated psilocybin dosing.[25]

While differences in dosing strategies, species, strains, and outcome measures certainly account for some of this variability in the observed outcomes of pre-clinical studies to date, the experimental model chosen also likely has a substantial impact. Indeed, no universal animal model exists for MDD or anxiety disorders, and even clinically-validated antidepressants exhibit differing levels and time courses of activity across the various models available.[27] However, it has been suggested that the differential sensitivity of these models may be leveraged to allow for a more nuanced understanding of the conditions under which a given biological mechanism is most relevant for a given treatment approach.[28] For example, chronic oral (PO) corticosterone exposure is a commonly used model in rodents when investigating altered hypothalamic-pituitary-adrenal (HPA) axis function in the context of psychiatric disorders; although HPA axis dysfunction is variable across clinical psychiatric populations, this approach generates HPA axis suppression similar to that found in individuals with acute and severe remitting MDD.[27] It also decreases hippocampal concentrations of brain-derived neurotrophic factor (BDNF), which has been suggested to be a supportive element for the aforementioned pro-neuroplastic effects of classical psychedelics observed in rodents.[29] Furthermore, this model has been successfully employed to distinguish between classical antidepressants with delayed onset of efficacy and rapidly-acting antidepressant compounds, and permits assessment of their potential utility in treatment-resistant depression.[30]

Building from this established literature, the chronic oral corticosterone exposure model appears to be relevant for the assessment of mechanisms supporting psilocybin’s variable subjective effects, which in humans can range from bliss to dread.[31] Importantly, both anxious effects and plasma cortisol concentrations have been reported to be transiently elevated by high-dose psilocybin treatment in healthy human subjects.[32] Furthermore, a particular form of acute anxiety, ‘dread of ego dissolution’, has been negatively associated with therapeutic efficacy in individuals with treatment resistant depression.[33] However, the causal relationship between acute cortisol release, anxiety, and alterations of therapeutic efficacy has not been directly assessed. Elevated plasma cortisol in humans may be directly leading to manifestation of anxious behavior, it may exclusively be acting as a downstream biomarker of an independently initiated ‘dread of ego dissolution’ psychological process, or there may be mutually-reinforcing feedback ongoing between these physiological and psychological processes. Furthermore, acute cortisol release and acute anxiety may each be correlated with worse therapeutic effects without both, or either, having to be causally related to this outcome. Indeed, observational study has identified a positive correlation between acutely challenging psychedelic experiences and long-term well-being, and at least one model of psychedelic efficacy proposes that overcoming difficult experiences through acceptance is of therapeutic benefit overall.[34,35] Interestingly, this model is consistent with stress inoculation effects seen in pre-clinical studies of anxious responding.[36,37]

As the deliberate induction of challenging experiences through increases in environmental stressors or other manipulations are ethically impermissible in psychedelic clinical trials, this study proposes use of rodent models as an important means by which to potentially help resolve this issue. The well-validated chronic PO corticosterone exposure model can be used to suppress rodent HPA axis function, while agonism of 5-HT_2A_R by psilocybin seems likely to activate HPA axis function and promote acute glucocorticoid release, as has been observed for other tool compounds in this class.[38] Using these primary manipulations, this work tests the hypothesis that chronic PO corticosterone exposure will decrease anxious responsiveness in mice treated with psilocybin at acute (0 - 1 h), post-acute (4 – 24 h) and long-term (2 - 8 days) time periods, due to suppression of drug-induced glucocorticoid release.

## 2. Materials and Methods

### 2.1 Animals and Husbandry

All mice used in this work were acclimated to University of Wisconsin vivarium conditions for at least seven-days prior to handling or experimentation. Food pellets (LabDiet) and water (Inno-Vive) were available *ad libitum,* unless otherwise noted. All C57Bl6/J mice (male; 6-8 weeks old; The Jackson Laboratory, ME, USA) were housed in groups of three or four, while under a 12hr artificial light/dark cycle. Room temperature remained constant between 22 – 24 °C. All experimental procedures were approved by the University of Wisconsin, Madison Animal Care and Use Committee (IACUC) and completed in full accordance with Research Animal Resources and Compliance (RARC) guidelines.

### 2.2 Drugs

All controlled substances were handled by authorized users on the Schedule I, Schedule II-V DEA research licenses, and WI Special Use Authorizations held by Dr. Cody Wenthur. See Supplementary Materials and Methods for details on sources, preparation, and administration.

### 2.3 Behavioral Tasks

To measure acute (0 – 1 h post-injection) unconditioned responses to drugs, mice were assessed for locomotor behavior in an open field test (OFT) and for perceptual alteration using an automated head twitch response (HTR) detection protocol adapted from previous approaches.[39,40] To measure post-acute (4 – 24 h post-injection) behavioral responses to drugs, mice were assessed in independent tests, including a forced swim test (FST), sucrose preference test (SPT), novelty suppressed feeding (NSF) and measurement of time spent in the center of the OFT. For assessment of drug effect sensitivity to corticosterone treatment, animals were tested in independent experimental batteries incorporating the OFT, SPT, and NSF tasks. See Supplementary Materials and Methods for details of the conditions used for each of these tests.

### 2.4 Corticosterone ELISA

Animals were anesthetized with isoflurane and trunk blood was collected into microcentrifuge tubes following decapitation. The samples were then centrifuged at 10,000 RPM for 10 min, the plasma fraction was separated, and stored at −80°C. Upon thawing, the plasma corticosterone concentrations were assessed using a colorimetric ELISA analysis (Enzo-Life Sciences, Corticosterone ELISA Kit) per the enclosed protocol. All independent biological samples were run in technical duplicate or triplicate.

### 2.5 Statistical Analyses

Statistical analyses were performed using GraphPad Prism, version 9 (San Diego, CA). All tests were run as two-tailed analyses, setting p <0.05 as the threshold for significance. Data analyzed across time were assessed using paired-analysis approaches; all samples were otherwise considered to be independent for analysis purposes. One and Two-way ANOVA analyses, as well as their non-parametric analogues, were corrected for multiple comparisons when assessing differences across each condition. In all cases, ‘n’ refers to the number of individual animal subjects assessed for a given data point.

## 3. Results

In order to first identify a dose of psilocybin that could reliably demonstrate acutely psychoactive effects, 6-8 week old, male, C57Bl/6 mice were given IP injections with 3 mg/kg psilocybin, and observed for the induction of head-twitch responses (HTR), which are a classical proxy of hallucinogenic activity in mice (Figure 1A). Head twitches were detected automatically via fluctuations in magnetic field strength signals from magnets cemented to the animal’s skulls, using slight modifications from previously reported methods.[39,40] Additionally, a 30 mg/kg IP dose of ketamine was administered as a dissociative control that has rapidly acting antidepressant effects, but does not alter HTR (Figure S1A). Psilocybin-treated animals demonstrated a clear elevation in the number of head-twitch responses occurring in the 10 min after IP administration as compared to saline or ketamine (Two-way Repeated Measures (RM) ANOVA, F_interaction_ (36, 360) = 3.97; p <0.0001; Tukey’s, q (7.17) = 6.92; p = 0.0041 vs saline; q (6.36) = 7.33, p = 0.0042 vs ketamine). These head twitch responses rapidly returned to baseline during the 2 h post-injection measurement period. With this evidence of meaningful *in vivo* pharmacologic activity occurring acutely at 3 mg/kg of IP psilocybin, we next looked at effects of this intervention at the 4 h post-acute time period where the acute locomotor and behavioral disruptions resulting from this dose had subsided (Figure S2). At this post-acute time period, no psilocybin-induced changes in behavioral despair were observed, using immobility time in the FST (Figure 1B). In contrast, ketamine had robust effects on this outcome (Figure S1B, ANOVA, F (2, 31) = 30.56; p <0.0001; Sidaks; t (31) = 7.03, p <0.0001 vs. saline; t (31) = 6.36, p <0.0001 vs psilocybin). There was also no significant effect of psilocybin on hedonic responding, as measured by the sucrose preference post-acutely (Figure 1C). However, psilocybin treatment did demonstrate post-acute anxiolytic-like effects through reduction in the latency to first consumption in the NSF at 4 h post-treatment (Figure 1D; Student’s t-test; t (18) = 2.25; p = 0.0372).

**Figure 1.**
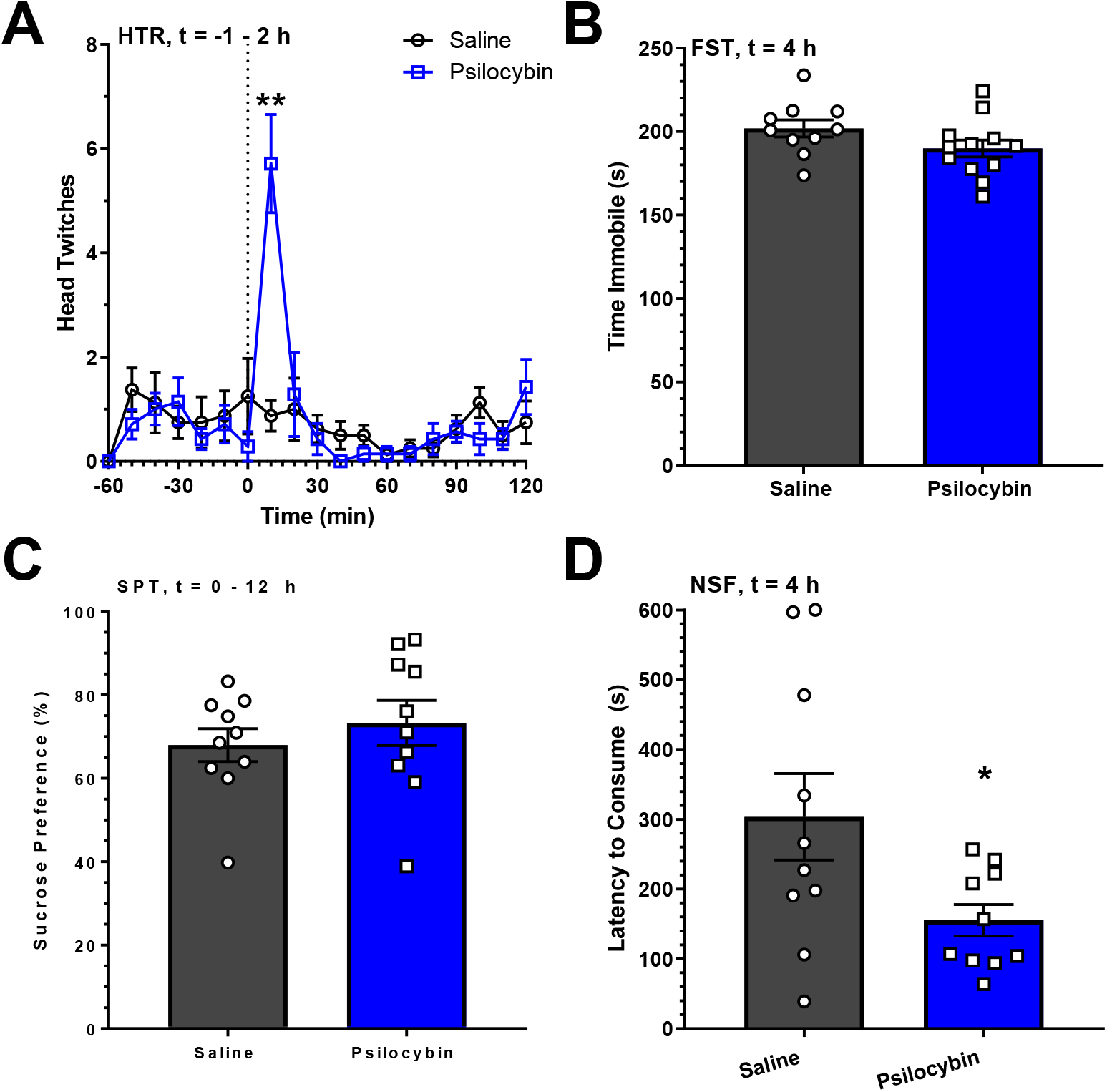
Psilocybin Has Post-Acute Anxiolytic Effects on Novelty Suppressed Feeding. A) Time course of automated head twitch response detections across each 10 min time period from 1 h before to 2 h after drug administration. (n=8, saline; n=7, psilocybin) **: p < 0.01. B) Immobility time in the forced swim test at 4 h following drug administration. C) Sucrose preference from 0 – 12 h after drug administration. D) Latency to feed in the novelty suppressed feeding test at 4 h after drug administration. * = p < 0.05. All experiments use 3 mg/kg psilocybin IP. All data presented as mean ± SEM.

To follow-up on these apparent psilocybin-induced post-acute anxiolytic effects, a dose response curve from 0 – 3 mg/kg was generated using total distance traveled and time spent in the center in the OFT. The relationship between acute and post-acute responding was assessed through measurement of these behaviors in two independent groups of animals; the first was assessed at the time of maximal HTR response (5 −15 min after injection), and the second was measured at the time of post-acute anxiolysis in the NSF (4 h). In regard to total distance traveled at 15 min (Figure 2A), there was a dose-dependent increase in activity (ANOVA, F (3,40) = 4.62; p = 0.0073); Post-hoc test for linear trend, F (1, 40) = 13.39; p = 0.0007) with 3 mg/kg of IP psilocybin showing a significant increase as compared to saline (Tukey’s, q (40) = 5.13; p = 0.0042). By 4 h after treatment, this overt locomotor effect had been lost (Figure 2B), and there was no significant correlation between locomotor activity at these two time points (Figure 2C). At 15 minutes (Figure 2D), low-dose (0.3 mg/kg) psilocybin increased the time in center of the field, while higher doses exhibited a dose-dependent reduction in this measure (ANOVA, F (2,27); p = 0.0137; Test for linear trend, F (1, 27) = 10.08; p = 0.0037). Intriguingly, at 4 h after treatment (Figure 2E), low-dose psilocybin (0.3 mg/kg) decreased the time in center, while higher doses exhibited a dose-dependent increase (ANOVA, F (2,26) = 3.37; p = 0.0498; Test for linear trend, F (1,26) = 6.72; p = 0.0155). The responses at the 15 min and 4 h time points were found to be anti-correlated across the dose-range tested (Figure 2F; Pearson’s correlation, R_2_: −0.95, p = 0.0264). This result indicates that psilocybin-induced acute anxiogenic responses are associated with delayed anxiolytic effects.

**Figure 2.**
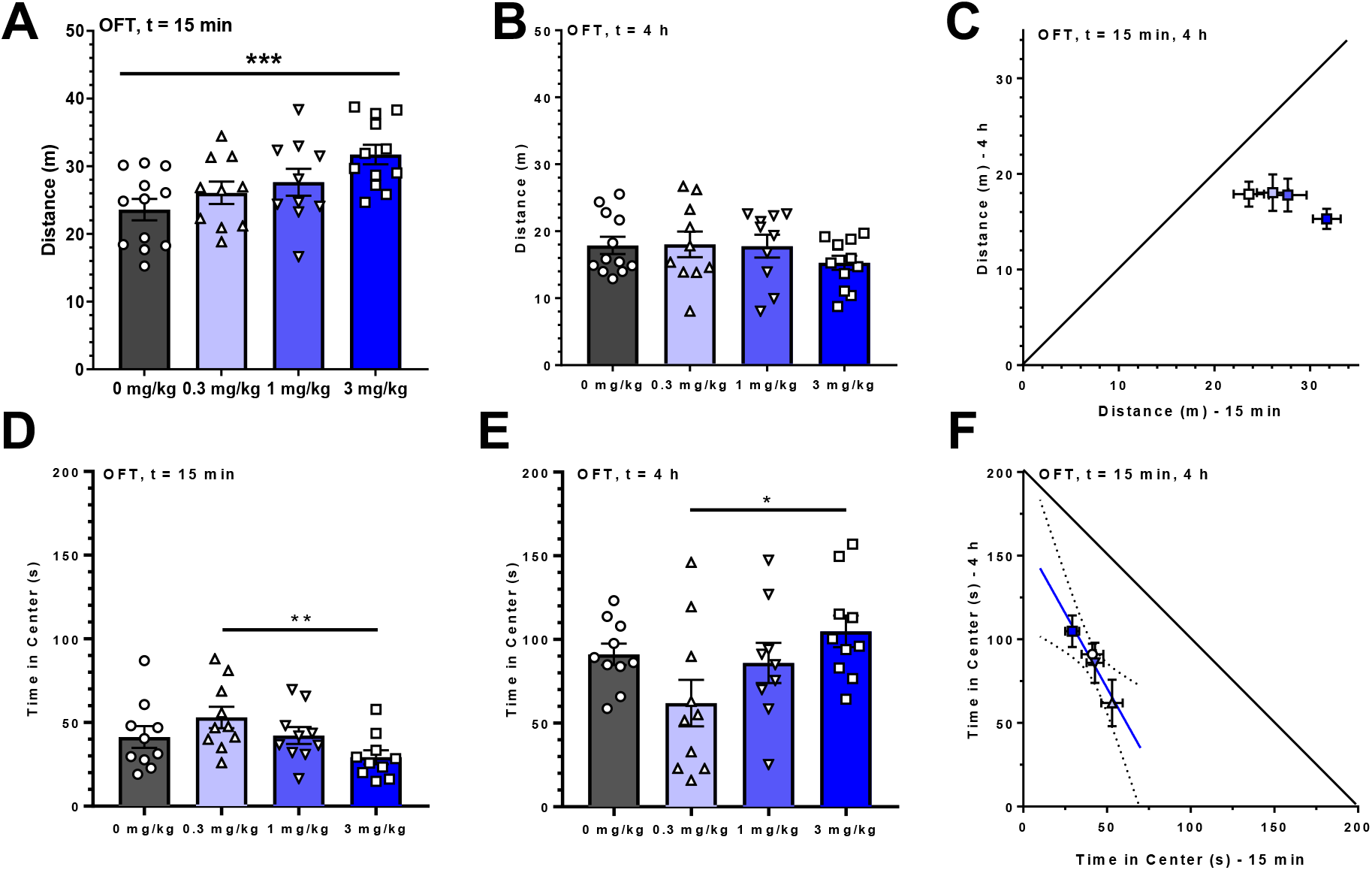
Psilocybin Dose-Dependently Increases Acute Anxiety-Like and Delayed Anxiolytic-Like Behaviors. A) Changes in locomotor response to increasing psilocybin doses in a 10 min open field test occurring from 5 – 15 min post-injection. ***: p < 0.001, ANOVA, test for trend. B) Changes in locomotor response to increasing psilocybin doses in a 10 min open field test beginning at 4 h post-injection. C) Correlative plot of distance traveled during the 15 min and 4 h open field tests. D) Time spent in the center of the arena in response to increasing psilocybin doses in a 10 min open field test occurring from 5 – 15 min post-injection. **: p < 0.001, ANOVA, test for trend. E) Time spent in the center of the arena in response to increasing psilocybin doses in a 10 min open field test beginning at 4 h post-injection. *: p < 0.05, ANOVA, test for trend. C) Correlative plot of time spent in the center of the arena during the 15 min and 4 h open field tests with linear regression and 95% CI. Data presented as mean ± SEM. For C and F, because different animals were assessed at 15 min and 4 h, the plot depicts the correlation between the means for each psilocybin dose, with the errors for these means added in quadrature.

With this indication that psilocybin’s acutely stressful actions are important for modification of post-acute responding, we next wanted to directly measure the effect of drug administration on acute, post-acute, and long-term glucocorticoid release. This outcome was measured in the presence of either no corticosterone, acute IP corticosterone pre-treatment, or chronic corticosterone PO exposure (Figure 3A). In the absence of any prior corticosterone treatment, plasma corticosterone concentrations were measured at 15 min, 4h and 7 days following IP injection with saline, 30 mg/kg ketamine (Figure S1C) or 3 mg/kg psilocybin (Figure 3B). The day 7 measurement occurred immediately following an OFT exposure to assess the impact of a non-drug stressor on subsequent corticosteroid release, which was used as a reference for the magnitude of effects for exogenous corticosterone administration below. While IP injection itself induced a transient elevation in plasma corticosterone (ANOVA, F (3, 42) = 32.49; p <0.0001; Tukey’s, q (42) = 6.786; p = 0.0001, saline vs baseline), both ketamine (Tukey’s, q (42) = 3.94; p = 0.0385) and psilocybin (Tukey’s, q (42) = 4.76; p =0.0086) induced a significantly greater increase than saline at 15 min, indicating that drug-dependent corticosterone release was also occurring. All groups had returned to baseline by 4 h post-treatment. No significant long-term differences between groups were observed following the delayed OFT exposure, indicating that the drug-treatment itself was not modifying subsequent stress-induced corticosterone release.

**Figure 3.**
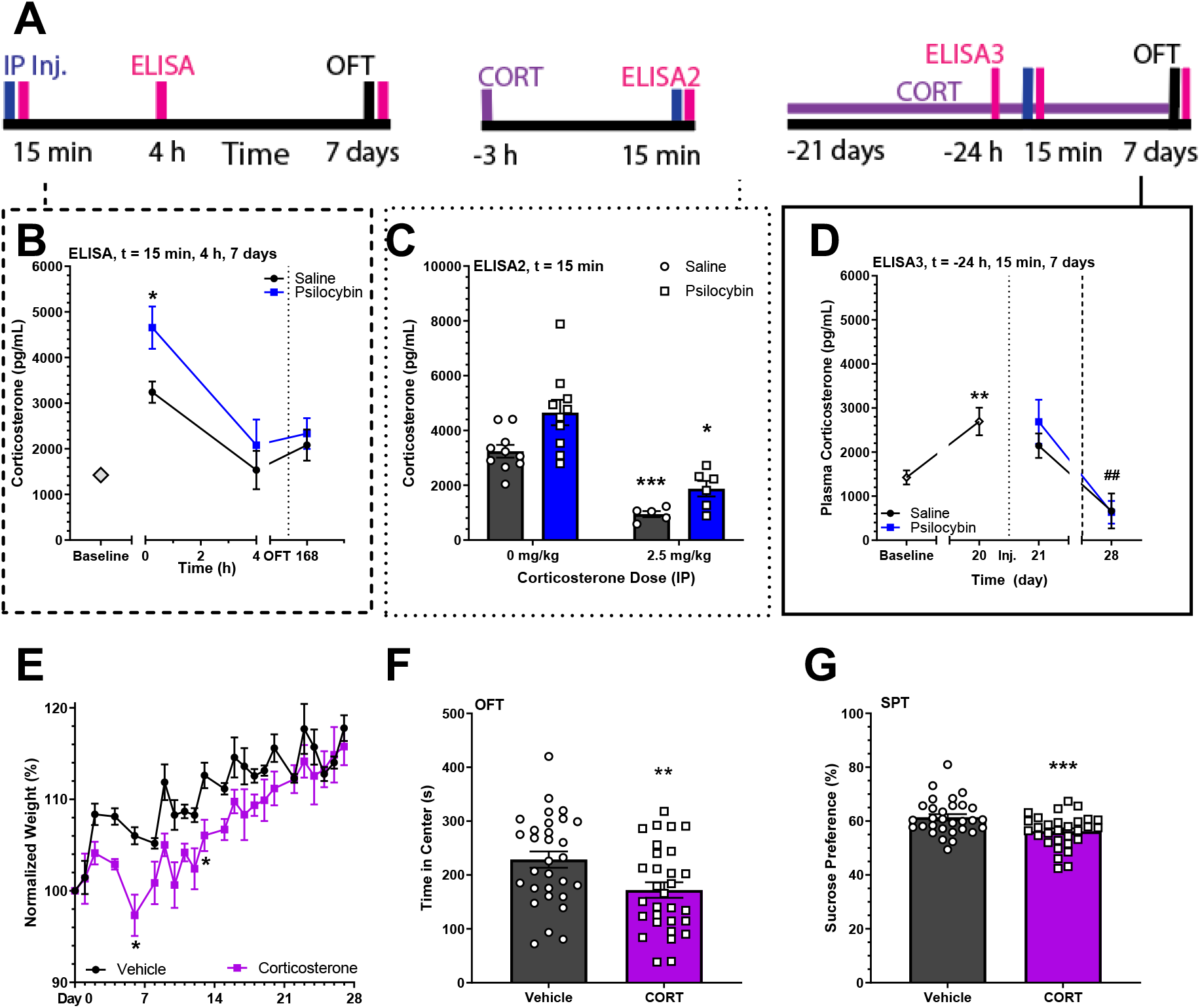
Chronic Corticosterone Exposure Suppress Stress and Psilocybin-Induced Acute Glucocorticoid Release. A) Experimental timelines for measurement of plasma corticosterone. IP Inj: intraperitoneal injection, ELISA: enzyme linked immunosorbent assay, OFT: Open Field Test; CORT: corticosterone. Vertical lines indicate acute intervention; horizontal lines indicate ongoing exposure. B) ELISA for plasma corticosterone concentrations at baseline and at various time-points following drug administration. Dotted line denotes open field test. Two-Way ANOVA with Sidak’s, * = p < 0.05 vs. Saline, 15 min. (n = 16 baseline, n = 10, saline, psilocybin). C) ELISA for plasma corticosterone concentrations at 15 min following drug administration in the presence of IP corticosterone pretreatment. Two-Way ANOVA with Sidak’s, * = p <0.05, *** = p <0.001 vs Saline + 0 mg/kg Corticosterone. D) ELISA for plasma corticosterone concentrations at baseline, following chronic oral corticosterone exposure (80 μg/mL), at 15 min following drug administration, and following exogenous corticosterone withdrawal. Student’s t-test, ** = p < 0.01 vs Baseline. ## = p <0.01 vs Vehicle. Dotted line denotes drug injection. (n = 16 baseline, day 20; n = 8, day 21 and 28). E) Normalized weight during oral corticosterone exposure. (n = 8) * = p <0.05 vs. Vehicle. F) Time spent in the center of an open field arena following 28 days of oral corticosterone exposure. ** = p < 0.01. G) Sucrose preference following 28 days of oral corticosterone exposure. *** = p < 0.001. All experiments use 3 mg/kg psilocybin IP. All data presented as mean ± SEM.

With this evidence that psilocybin-induced acute glucocorticoid release was occurring, we next endeavored to establish an effective exogenous corticosterone administration protocol for suppressing this response. Adapting an established rat protocol, acute corticosterone pretreatment at 2.5 mg/kg IP was first attempted (Figure 3C).[41] This acute intervention was able to significantly blunt all administration-associated glucocorticoid release at 15 minutes (Two-Way ANOVA, F_Cort_ (1, 41) = 76.71; p <0.0001). Blunting was also apparent for the pairwise comparisons between corticosterone pre-treated and saline-treated animals (Sidaks, t (41) = 4.43; p = 0.0003, saline; t (41) = 2.80; p = 0.0382, psilocybin; t (41) = 2.70; p = 0.049, ketamine; Figure S1D), but an overall drug effect was still measureable (Two-Way ANOVA, F_Drug_ (2, 41) = 6.515; p = 0.0035).

Passive exposure to 80 μg/mL of chronic PO corticosterone in drinking water for 21 days was next explored as a means to both establish HPA axis suppression, and as a chronic stress model relevant to induction of depressive-like and anxiety-like phenotypes. Use of this approach led to a significant increase in plasma corticosterone (Figure 3D; Paired t-test, t (15) = 3.17; p = 0.0063). Notably, the chronically elevated plasma corticosterone concentration seen in this model was similar to that transiently achieved following presentation of an OFT environmental stressor. Subsequent IP injection of psilocybin was unable to trigger an acute release of corticosterone. Furthermore, ongoing HPA axis suppression was assessed using plasma corticosterone concentrations on day 28; measurements occurred 7 hours after exogenous corticosterone withdrawal and 15 min after an OFT exposure to assess endogenous corticosteroid production and release in response to an environmental stressor. Indeed, HPA axis suppression was confirmed, as all chronic corticosterone-exposed mice had significantly lower concentrations of plasma corticosterone under these conditions (Student’s t-test; t (30) = 2.78; p = 0.0093).

In addition to these changes in acute glucocorticoid release and suppressed HPA axis function, the effects of chronic PO corticosterone exposure were also assessed in regard to metabolic and behavioral outcomes. Overall, animals chronically exposed to PO corticosterone demonstrated alterations in weight change over time (Figure 3E; Two-Way ANOVA: F_Cort_ (1, 375) = 53.03; p < 0.0001), a significant reduction in the time spent in the center of an open field, (Figure 3F; Student’s t-test, t (58) = 2.71; p = 0.0088), and a significant reduction in sucrose preference (Figure 3G; Student’s t-test, t (58) = 3.18; p = 0.0024) as compared to vehicle exposed animals. Overall, this indicated that the chronic corticosterone model was itself acting to enhance the expression of anxious and anhedonic behaviors.

Next, to assess the impact of psilocybin treatment on anxious and hedonic responding, animals chronically exposed to either PO corticosterone or vehicle were treated with either saline or 3 mg/kg psilocybin IP and tested in experimental batteries that incorporated longitudinal SPT, post-acute NSF, and delayed OFT exposures (Figure 4A). Together, these batteries were designed to assess drug effects on depressive-like and anxiety-like responding across multiple behavioral measures. Additionally, the order and timing of task presentation was varied in relationship to corticosterone exposure in order to differentiate the impact of blocking psilocybin-induced glucocorticoid release from the impact of altering the steady-state corticosterone concentrations surrounding the time of drug administration.

**Figure 4.**
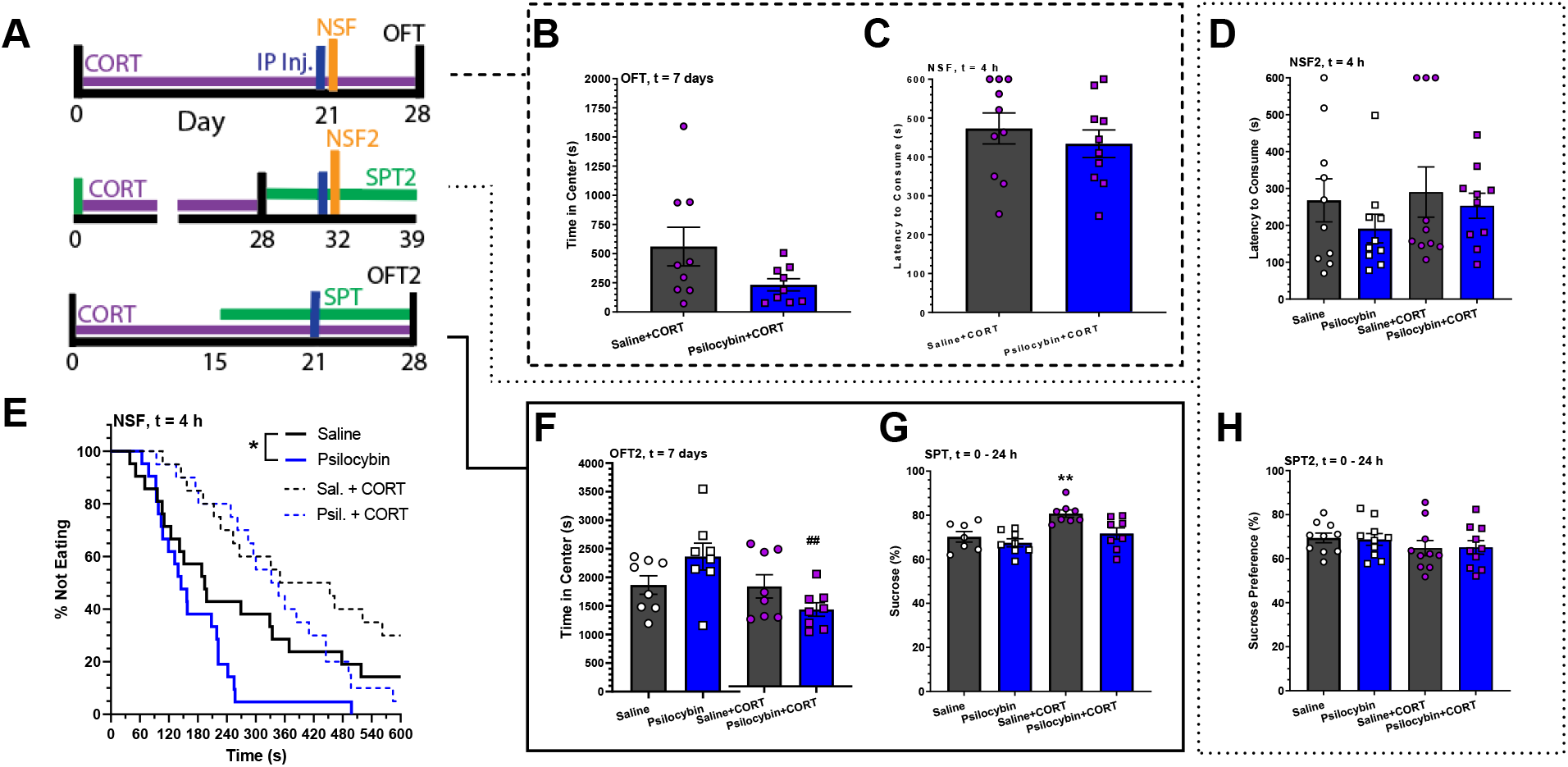
Chronic Corticosterone Exposure Blunts or Reverses Psilocybin’s Post-Acute and Long-Term Anxiolytic Effects. A) Experimental timelines for measuring the behavioral impacts of chronic oral corticosterone exposure on psilocybin treatment. IP Inj: intraperitoneal injection, NSF; Novelty Suppressed Feeding Test, OFT: Open Field Test; CORT: corticosterone. Vertical lines indicate acute intervention; horizontal lines indicate ongoing exposure. B) Time spent in the center during a 150 min open field test following oral corticosterone exposure at 7 days after psilocybin treatment. C) Latency to feed in the novelty suppressed feeding test during ongoing oral corticosterone exposure at 4 h after drug administration. D) Latency to feed in the novelty suppressed feeding test following prior oral corticosterone exposure at 4 h after drug administration. E) Survival curves of latency to feed for saline and psilocybin as matched across corticosterone exposure conditions (n = 20) *: p <0.05 F) Time spent in the center during a 150 min open field test following oral corticosterone or vehicle exposure at 7 days after psilocybin treatment. ## = p <0.01 vs. psilocybin. G) Sucrose preference during ongoing vehicle or corticosterone-exposure at 0 – 24 h after psilocybin administration. #: p < 0.05, ###: p < 0.001 vs. Saline + CORT H) Sucrose preference following vehicle or corticosterone-exposure at 0 – 24 h after psilocybin administration. **: p < 0.01 vs. saline. All experiments use 3 mg/kg psilocybin IP. All data presented as mean ± SEM.

When animals were assessed in the OFT following 28 days of PO corticosterone exposure and 7 days after IP drug administration, psilocybin treatment resulted in a non-significant decrease in center time at this remote time point (Figure 4B). This contrasted with the delayed anxiolysis previously seen following psilocybin treatment alone. Likewise, when the NSF was administered on day 21, d psilocybin’s previously-observed reduction in the latency to feed was absent; animals exhibited behavior that was no different from saline treatment (Figure 4C). Notably, elimination of both of these behaviors by chronic corticosterone administration occurred despite the fact that they were administered under different steady-state conditions in regard to plasma corticosterone; the OFT was administered after withdrawal of exogenous corticosterone to yield a suppressed baseline, and the NSF was administered during an ongoing period to yield an elevated baseline.

Indeed, the NSF was further presented to a separate group of animals during a period of steady-state glucocorticoid suppression to address the idea that acute suppression of drug-induced glucocorticoid release was more relevant to this loss of psilocybin-effect than the steady-state alteration for corticosterone. As before, the effect of psilocybin in the NSF was diminished (Figure 4D). Interestingly, this was also true for vehicle treated animals, implying that environmental stressors such as the single-housing conditions used during this battery may still be relevant modifiers of response when acute glucocorticoid release is intact. To directly assess the impact of corticosterone pretreatment itself, psilocybin’s effects on latency to feed in the NSF were then assessed across all naïve/ vehicle conditions vs all chronic corticosterone conditions (Figure 4E). A direct comparison of the psilocybin and saline profiles for naïve/ vehicle-treated animals demonstrated a significant anxiolytic effect of psilocybin (Log-rank (Mantel Cox) test; χ_2_ (1) = 4.074; p = 0.0436; Hazard Ratio = 0.50, 95%CI [0.25 – 0.98], while this anxiolysis was absent when chronic corticosterone exposure had occurred. No effect of ketamine was observed at the dissociative 30 mg/kg dose (Figure S1E).

In light of psilocybin’s anxiogenic-like response seen in the OFT following chronic corticosterone exposure, another behavioral battery was run in order to more fully determine the effects of suppressing acute corticosteroid release on long-term measures of psilocybin response. In this experiment, another group of animals were again passively exposed to either vehicle or chronic corticosterone PO for 28 days and administered 3 mg/kg IP psilocybin on day 21 of exposure. These mice were then longitudinally assessed in the SPT, followed by assessment in the OFT on day 28. In regard to time spent in the center of the OFT at this 1-week time point after psilocybin administration (Figure 4F), a clear interaction between psilocybin and corticosterone effects was observed (Two-Way ANOVA: F_interaction_, (1,28) = 5.95; p = 0.0213). Animals exposed to chronic corticosterone and treated with psilocybin spent significantly less time in the center of the arena than unexposed animals treated with psilocybin (Sidak’s, t (28) = 3.605; p = 0.0072).

Notably, the two delayed OFT experiments in these batteries were undertaken during opposing phases of the animals’ light/dark cycle; corticosterone concentrations fluctuate significantly across these periods.[42] Nevertheless, anxiety promoting effects of corticosterone pretreatment were observed in both conditions (Figure S3). This observation is again supportive of the idea that suppression of transient, psilocybin-induced glucocorticoid release is the behaviorally relevant outcome arising from chronic corticosterone exposure, and that alterations in steady-state glucocorticoid concentrations overall have little impact when such suppression is in effect, even for behavioral anxiety measurements occurring at remote time points.

In contrast to the OFT results, the most notable effects of corticosterone exposure on psilocybin effects in the SPT occurred acutely. Specifically, the outcome profile was consistent with psilocybin acutely masking a compensatory adaptation to avoid PO corticosterone alone (Figure S4, Figure 4G; ANOVA, F (3, 27) = 7.49; p = 0.0008; Sidak’s, t (27) = 3.441; p = 0.0057 saline vs saline+CORT). This lack of direct psilocybin (or ketamine (Figure S1F)) effects on sucrose preference was confirmed post-acutely in animals exposed to an SPT that occurred after PO corticosterone was no longer available in their water (Figure 4H). No long-term effects of psilocybin or ketamine on sucrose preference were observed in the week following administration under any of the conditions tested in these batteries (Figure S5).

Overall, these observed effects of chronic corticosterone exposure on alteration of post-acute and long-term anxiolytic-like responses of psilocybin, indicate that HPA axis function and drug-induced glucocorticoid release are able to influence the delayed effects of this drug. However, given that this acute corticosteroid release is occurring alongside perceptual disturbances and induction of other relevant signaling cascades, we decided to assess whether acute glucocorticoid release alone was sufficient to alter delayed anxiolytic-like responding. Here, the OFT results from those animals used for ELISA analysis of acute, post-acute, and delayed plasma corticosterone concentrations were assessed, since these animals had directly validated acute release of corticosterone. These animals differed from other OFT exposed animals in a critical way, however – in order to collect the plasma samples for the ELISA, each mouse had been exposed to brief (3-5 min) isoflurane anesthesia at 15 minutes after psilocybin administration. For these animals, no long-term effects of psilocybin were observed either in the absence (Figure 5A) or presence (Figure 5B) of prior chronic PO corticosterone exposure. Correspondingly, in contrast to the reversal of psilocybin-dependent effects in the OFT that was seen in animals not exposed to isoflurane (Figure 5C; Student’s t-test, t (14) = 2.607; p = 0.0207), no interactions were seen in the delayed OFT for these animals that had been exposed to isoflurane anesthesia shortly after psilocybin administration (Figure 5D). This indicates that acute glucocorticoid release alone is not sufficient to induce delayed anxiolytic-like effects.

**Figure 5.**
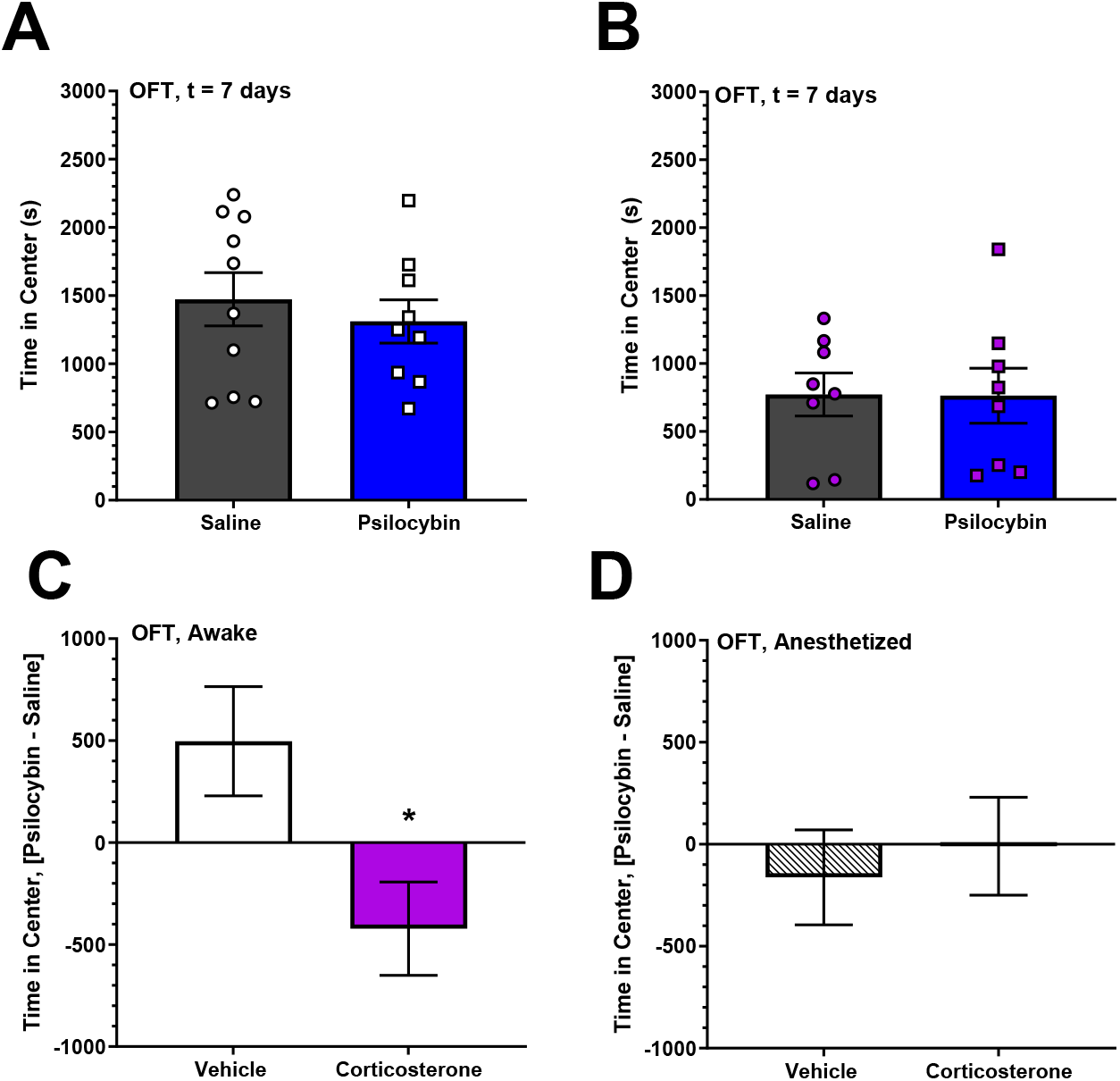
Brief Isoflurane Anesthesia Eliminates the Interaction Between Corticosterone Exposure and Psilocybin’s Long-Term Effects on Anxious Responding. A) Time spent in the center during a 150 min open field test at 7 days after psilocybin treatment. B) F) Time spent in the center during a 150 min open field test following oral corticosterone or vehicle exposure at 7 days after psilocybin treatment. C) Difference between psilocybin mean response and saline mean response for time in center during a 150 min open field test at 7 days after psilocybin treatment. * = p < 0.05. D) Difference between psilocybin mean response and saline mean response for time in center during a 150 min open field test at 7 days after psilocybin treatment., in animals exposed to brief (3-5 min) isoflurane anesthesia at 15 min after psilocybin treatment. All experiments use 3 mg/kg psilocybin IP. Data presented as Mean ± SEM. For C and D, this represents the difference between the means of the psilocybin and saline treated animals, with the errors for these means added in quadrature.

## 4. Discussion

In these experiments, it was found that psilocybin can induce post-acute (Figure 1D, 2E) anxiolytic-like effects. Such post-acute outcomes were seen to be correlated with acute anxiogenic effects (Figure 1E). Chronic corticosterone exposure was determined to eliminate psilocybin-induced corticosterone release occurring during the acute anxiogenic period (Figure 3D), to block post-acute psilocybin anxiolysis (Figure 4E) and to reverse its long-term effects from anxiolytic-like to anxiogenic-like (Figure 5C). Surprisingly, these outcomes support the conclusion that the delayed anxiolytic-like behavioral effects of psilocybin in male mice are supported by, but not exclusively a result of (Figure 5D), drug-induced glucocorticoid release occurring at the time of drug administration. The interaction between glucocorticoid manipulation and behavioral effect was most pronounced in the time in center OFT results measured one week after psilocybin administration, but a reduction in psilocybin’s anxiolytic effects on feeding latency in the NSF was also observed following chronic corticosterone exposure. A notable limitation of these results is the use of male mice only; explicit assessment of sex differences in this response should absolutely be pursued in future work, particularly given well-established knowledge regarding differential stress responses in male and female mice.[43,44] Another limitation is the inability to fully eliminate the role of IP injection-related stress versus drug-related stress in the vehicle treated animals; while the dose-dependency of acute anxiogenic and delayed anxiolytic-like effects in the OFT is indicative of a drug-specific response, the use of minimally stressful administration techniques for psilocybin (such as infusion through an IV catheter) could lend further clarity to this issue in future studies.

To contextualize these findings, it is notable that repeated corticosterone administration alone has recently demonstrated bidirectional effects on stress-associated behavioral adaptation in rodents; a single corticosterone exposure can enhance future anxiety-like responding, but a rapid stress or corticosterone post-exposure is sufficient to prevent this behavioral change.[45,46] Psilocybin administration may be similarly acting as an initial stressor that provides resilience to subsequent stress. Furthermore, as the modifying effects of repeated corticosterone have been hypothesized to be dependent on delayed glutamate-sensitive neuroplastic effects occurring in the basolateral amygdala, future investigations into the mechanistic basis for the observed interaction between psilocybin and corticosterone will benefit from monitoring both immediate and long-term changes in cortical and amygdalar glutamate concentrations following 5-HT_2A_R agonist administration.[46,47]

The blunted NSF outcomes observed following psilocybin administration in the context of single- vs group-housing suggest that rodent models may be useful for addressing the functional impact of differing environmental conditions at the time of psychedelic drug onset. Crucially, such environmentally-dependent effects have already been observed regarding the effects of selective serotonin reuptake inhibitors in mice. [48] Nevertheless, it is worth highlighting that our identified interaction between acute glucocorticoid release and delayed anxiolytic effects was observed in a mammalian species that is not susceptible to the psychedelic messaging encoded within human socialization processes or subsequent development of expectation biases, and has limited evidence demonstrating self-awareness or narrative meta-cognition. [49,50] This outcome ultimately supports the suggestion that pharmacologically-mediated neuroplastic mechanisms comprise an important part of psychedelic drug effects. Isoflurane’s directly suppressive effects on hippocampal neuroplasticity may thus be especially relevant to the loss of psilocybin effects observed following anesthesia, although the potential influence of lost perceptual input during anesthesia cannot be ruled out.[51,52] Ultimately, further studies using explicit measurement of neuroplasticity and application of direct environmental manipulation are needed to address this translationally-relevant issue.[31,53–56]

Importantly, the delayed anxiolytic effects of psilocybin and its interaction with glucocorticoid release were not uniformly present across all tasks studied here. While some acute and post-acute effects on sucrose preference were observed following psilocybin administration, no long-term effects on hedonic response were seen. Additionally, we found no post-acute effects on immobility time in the FST.[25] However, these negative results may reflect specific limitations of the experimental parameters chosen here, rather than a failure of these models to recapitulate clinical findings with psilocybin treatment in general. For example, alternative mouse strains may be more appropriate for SPT measurements, as C57BL/6J mice demonstrate relatively low sensitivity to chronic corticosterone treatment in the SPT.[57] Additionally, the previous studies that have successfully observed psilocybin-mediated effects on FST in mice used more remote time-points (<7 days) for measurement, indicating that longer delay periods may result in a different outcome than observed in this work.[21]

In conclusion, these findings demonstrate that chronic corticosterone suppression of drug-induced acute glucocorticoid release is a beneficial model for the study of neuroendocrine impacts on psychedelic effects in rodents. Whether acute glucocorticoid release similarly supports delayed enhancement of stress-resilience in clinical trials using psilocybin remains an important question for translational consideration, particularly in concert with assessment of basal HPA axis dysfunction across different treatment populations. Given the limited mechanistic information available in this area thus far, numerous immediately relevant future directions are also available for rodent studies. These include assessment of whether the acute glucocorticoid-delayed anxiety interaction generalizes to other means of manipulating HPA axis function (including hyper-activation) and to other psychedelic drug classes in rodents, as well as measurement of functional and structural neuroplastic outcomes following coincident glucocorticoid release and 5-HT_2A_R activation, and dissociation of the role of anxiety-associated perceptual alterations and stress-modified neuroplastic changes in the observed isoflurane blockade of psilocybin’s long-term effects.

## Acknowledgements

This work was supported through grant funding to Cody J Wenthur from the National Institute of Mental Health (R01MH122742), fellowship for Nathan T Jones from the National Institute of General Medical Services (T32GM008688), fellowship for Zarmeen Zahid from the National Institute of Neurological Disorders and Stroke (T32NS105602), and internal funding from the University of Wisconsin – Madison Schools of Pharmacy / Medicine and Public Health to Cody J Wenthur and Matthew I Banks. Psilocybin for this study was provided to the investigators free-of-charge by the not-for-profit USONA Institute. The authors thank Robert Kargbo and Alex Sherwood of the USONA Institute for synthesis and helpful discussions related to the storage, stability, and analysis of psilocybin. The authors thank Clara Nickel for performing blinded analyses of the NSF videos.

## Author Contributions

*Nathan T Jones:* Investigation, Methodology, Formal Analysis, Visualization, Validation, and Writing – Review and Editing. *Zarmeen Zahid:* Investigation, Methodology, Formal Analysis, and Writing – Review and Editing. *Sean M Grady:* Investigation, Data Curation, Visualization, Writing – Review and Editing. *Ziyad W Sultan*: Investigation, Data Curation, Writing – Review and Editing. *Zhen Zheng:* Investigation, Validation, Writing – Review and Editing. *Matthew I Banks:* Conceptualization, Validation, Resources, Writing – Review and Editing, Supervision, Funding Acquisition. *Cody J Wenthur:* Conceptualization, Validation, Formal Analysis, Investigation, Resources, Writing – Original Draft, Visualization, Supervision, Project Administration, Funding Acquisition.

## Appendix A. Supplementary Methods and Materials

## Surgery

Skull screw EEG electrodes were chronically implanted in animals under isoflurane anesthesia (1.5 - 2%) using aseptic technique. Electrodes were placed bilaterally in the frontal (1.5 mm anterior to Bregma, 1.5 mm lateral to midline) and parietal (2.0 mm posterior to Bregma, 2.0 mm lateral to the midline) plates. Bilateral reference electrode screws were placed through the occipital plate and tied together to ground. EEG wires were stranded copper (0.012” diameter, 0.022” diameter including insulation; Cooner Wire, Chatsworth, CA). Wires were soldered to a 1-cm2 electrode interface board (EIB-16; Neuralynx, Bozeman, MT), which was fixed into place using dental cement (Fusio A3; Pentron; Orange, CA). Animals recovered for at least 5 days prior to the first recording day and were housed individually.

## Drug Preparation and Administration

Psilocybin powder (Usona Institute; Madison, WI) was diluted in 0.9% sterile saline, then acidified to a pH of 1-2 with 1 M HCl, sonicated for 30-60s, and brought to pH 6-7 using 1 M NaOH. This material was filtered through a 0.2 um filter, and administered intraperitonally (IP) at doses between 0.3 – 3 mg/kg. Ketamine Hydrochloride (Spectrum Chemical Mfg. Corp.; Gardena, CA) was diluted in 0.9% sterile saline, filtered through a 0.2 um filter, and administered at a dose of 30 mg/kg IP. All IP injections were given at a volume of 10 mL/kg. Corticosterone (Sigma-Aldrich) was diluted in 10% ethanol (EtOH) or 4.5% (2-hydroxypropyl)-Beta-Cyclodextrin (Biosynth-Carbosynth) in water, vortexed for 1 min, and then sonicated for 3 min at 22 °C, before being diluted to either 1% EtOH or 0.45% Beta-cyclodextrin in the animals’ drinking water. For experimental batteries requiring chronic corticosterone exposure, mice were given ad libitum access to either corticosterone water (0 - 80 μg/mL) or vehicle (1% EtOH or0.45% (2-hydroxypropyl)-Beta-Cyclodextrin) in their home cage for 28 days. [44–46]. Both vehicle and corticosterone bottles were refreshed every 7 days for the duration of the 28-day period.

## Open Field Test (OFT)

To test for drug-induced changes in locomotor, exploratory, and anxious behavior, mice were assessed in the OFT. Mice were injected IP with psilocybin (0 - 3 mg/kg) and then individually placed into a corner within an open-field apparatus (41×20×24 cm), at a time period from 5 min to 7 days afterward. The center zone was defined as the middle one-third (6×27cm) of the arena. The apparatus was illuminated at ~200 - 250 lux. Mice were allowed to explore freely for 10 minutes in acute experiments and 150 min in long-term experiments. Time spent in the center and total distance travelled were automatically quantified using the Any-Maze software. Each apparatus was cleaned before and after each test with Trifectant. All OFT measurements were run between 1-4 PM. For all stand-alone OFT experiments, this represented the dark phase of the cycle. In behavioral batteries where the OFT was given subsequent to the NSF, this represented the light phase of the cycle.

## Head Twitch Response (HTR)

To measure acute unconditioned responses to psilocybin, mice were assessed using an automated HTR detection platform adapted from previous approaches. [47,48] In this study, mice already implanted with chronic skull screw EEG electrodes were anesthetized with isoflurane (1.5 - 2%) to attach a neodymium magnet to the exposed dental cement from the EEG implant, at least 1 day prior to recording. Following recovery, individual animals were placed into a clear acrylic cylinder (15.24 cm height × 15.24 cm diameter) wrapped with ~300 rotations of 30 Gauge copper magnet wire, (Essex, Fort Wayne, IN) inside of a dark sound-attenuation chamber, connected to a flexible tether (ZC16) to record EEG while allowing free range of motion, and their behavior was recorded using an infrared camera (240×320 pixels) controlled by Synapse (Tucker Davis Technologies, Alachua, FL [TDT]) for 1 h prior to drug administration. Magnetometer signals were amplified near the source with a homemade custom circuit, and the signal was routed to an RZ5D (filtered at 0.2-1000 Hz, then digitized at 3,051.8 Hz). After this time period, the animals were administered 3 mg/kg IP psilocybin and recorded for an additional 4 h. Changes in the local magnetic field induced by head twitches (~ 60-90 Hz signal) were assessed using in-house MATLAB code. Automated results were compared to observed HTRs for internal validation. All HTR responses were recorded during the dark phase of the light cycle.

## Forced Swim Test (FST)

To measure the post-acute effects of drug treatment on immediate threat response, mice were assessed in the FST. Mice were injected IP with either psilocybin (3 mg/kg), ketamine (30 mg/kg) or saline. Animals were individually placed into a clear Plexiglas swim tank (46×10cm) for a period of 6 min. Water temperature was maintained at 26°C and the tanks were illuminated at 40-42 lux. Immobility time and distance travelled were quantified for the final 4 min of the test using Any-Maze software. At the completion of the testing period, animals were removed from the water, dried and placed into a clean cage with a heating pad to facilitate rapid recovery of normal body temperature. The animals were monitored in this chamber for 15-minutes before being placed back into group housing. All FST measurements were run between 1-4 PM, during the dark phase of the cycle.

## Sucrose Preference Test (SPT)

The SPT was conducted as a measure of hedonic responding both acutely and chronically. In this test, mice were placed into individual housing (41×20×24cm) to habituate for 48 h and then provided two identical bottles with either a 1% sucrose solution or water. At this time, mice had ad libitum access to standard chow and the 2 bottles provided. Animals underwent three 16 h restriction periods during which they had access to the 1% sucrose solution and water, with food and water provided between these periods. Immediately after the final restriction period, two identical bottles containing either 1% sucrose solution or water were placed into each cage and consumption was measured for a period of up to 15 days. The bottles were weighed at each test period, and sucrose preference was calculated as: sucrose weight / (sucrose weight + water weight) *100.

## Novelty Suppressed Feeding (NSF)

In order to measure animal behavior in a task combining motivated behavior with anxious responding, mice were assessed within the NSF. Animals underwent a sequential food reduction of 2-Days at 20% and 1-day at 80% and food deprivation (16 - 24 h,). Mice then received an IP injection of saline or drug 4-5 h prior to performing the NSF test. For the test, a food pellet soaked in 50% sucrose solution was placed into a glass petri dish that served as the feeding zone (9×0.375 cm) and centered within a novel cage environment (61×41×37cm) that was brightly illuminated. Mice were then placed into a corner of the apparatus and allowed to explore for 10 minutes. Latency to first feed was recorded by a trained and a blinded observer, and movement and distance traveled were monitored via the Any-Maze software. Pellet weights were also obtained immediately before and after each test. After testing, the mice were returned to their housing and given normal food and water. Each apparatus was cleaned before and after each test with Trifectant. All NSF measurements were run between 11-4 PM, during the light phase of the light cycle.

## Appendix B. Supplementary Data

**Figure S1.**
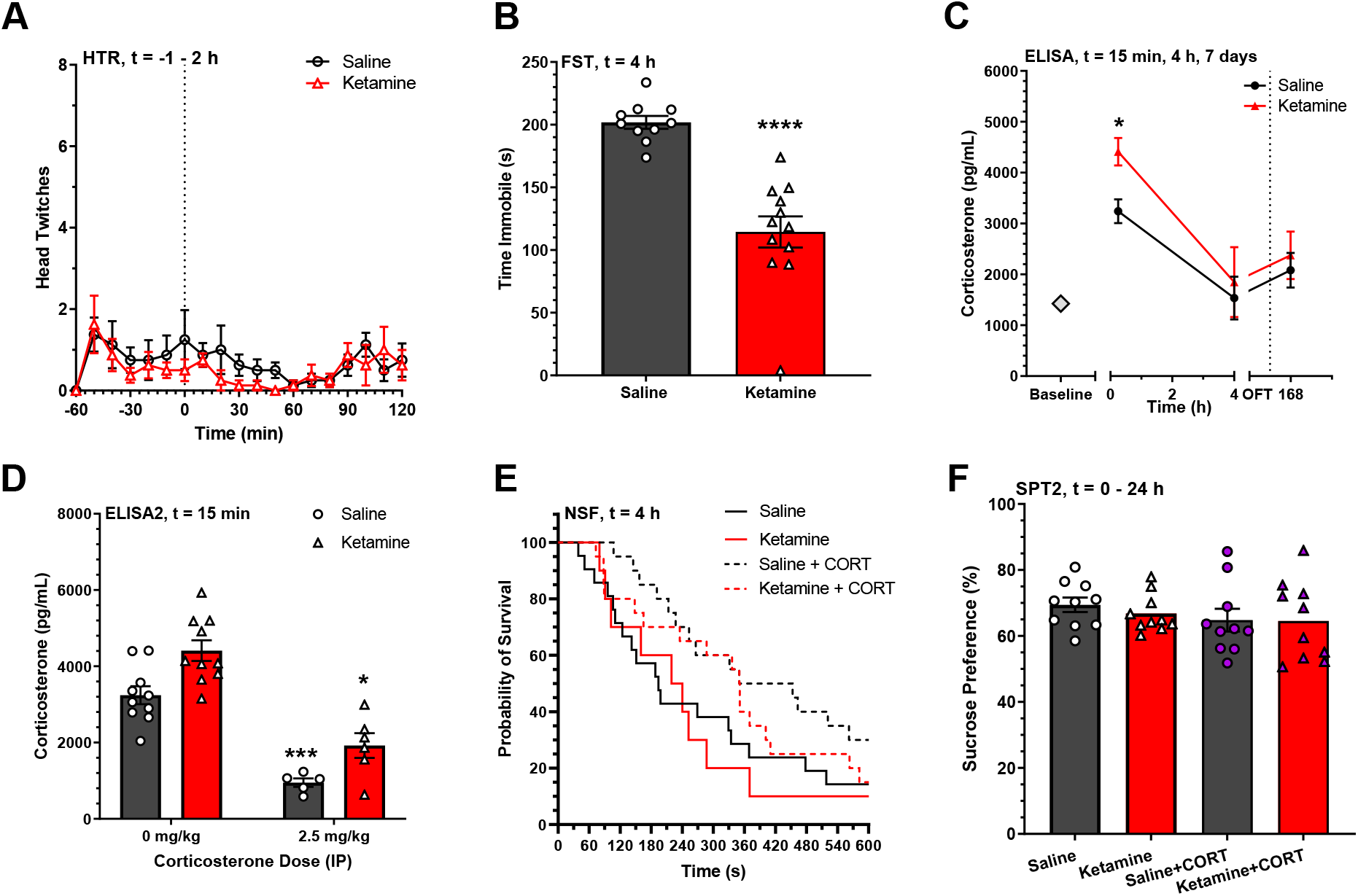
Effects of 30 mg/kg Ketamine on Study Measures. A) Time course of automated head twitch response detections across each 10 min time period from 1 h before to 2 h after drug administration. (n=8, saline, ketamine) B) Immobility time in the forced swim test at 4 h following drug administration. ****: p < 0.0001. C) ELISA for plasma corticosterone concentrations at baseline and at various time-points following drug administration. Dotted line denotes open field test. Two-Way ANOVA with Sidak’s, * = p < 0.05 vs. Saline, 15 min. (n = 16 baseline, n = 10, saline, ketamine). D) ELISA for plasma corticosterone concentrations at 15 min following drug administration in the presence of IP corticosterone pretreatment. Two-Way ANOVA with Sidak’s, * = p <0.05. E) Survival curves of latency to feed for saline and ketamine as matched across corticosterone exposure conditions (n = 10, ketamine; n = 20, all others). F) Sucrose preference following vehicle or corticosterone-exposure at 0 – 24 h after ketamine administration. All data presented as Mean ±SEM.

**Figure S2.**
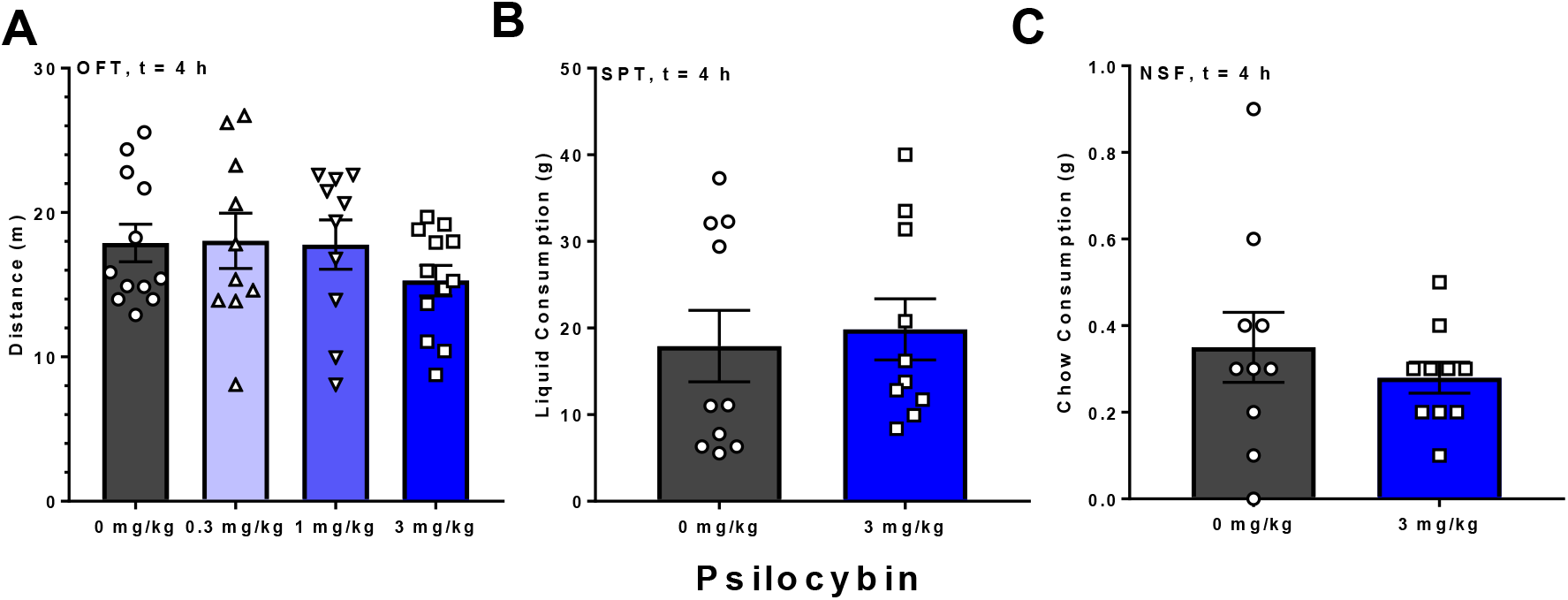
No Motoric or Consumptive Disturbances of 3 mg/kg Intraperitoneal Psilocybin Are Observed Post-Acutely at 4 Hours After Administration. A) Locomotor response to psilocybin doses in an open field test from 240-250 min after psilocybin injection. B) Total distance traveled in a forced swim test occurring 240 min after psilocybin injection. C) Immobility time from this forced swim test. D) Total liquid consumption (1% Sucrose and water) in a sucrose preference test for the period from 4 – 20 h after psilocybin injection. E) Sucrose preference from this sucrose preference test. F) Total chow consumption in a novelty suppressed feeding task for the period from 240-250 min after psilocybin injection. G) Latency to first consumption from this novelty suppressed feeding task. All error bars presented as mean ± SEM.

**Figure S3.**
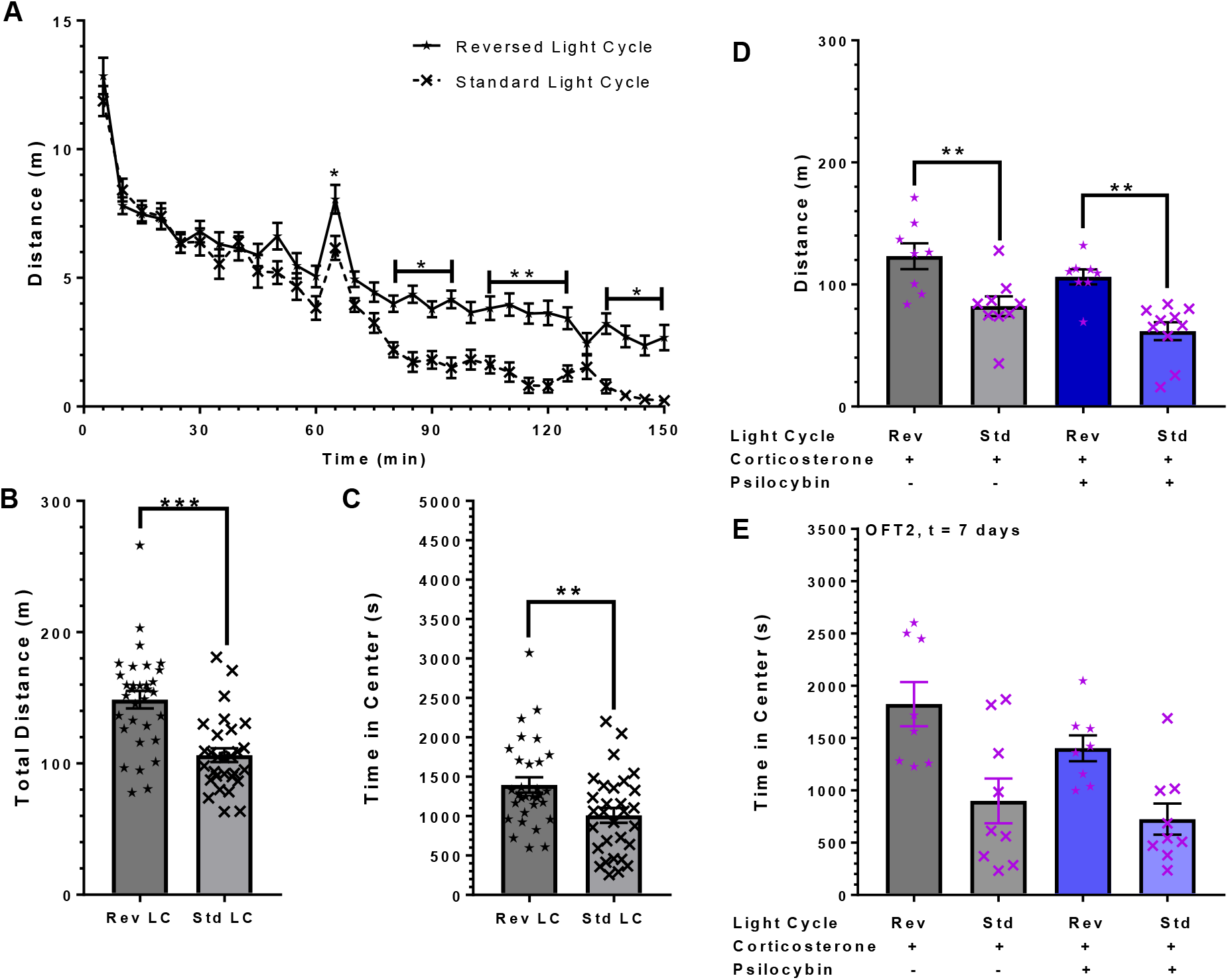
Light Cycle Reversal Increases Signal Window for Locomotor and Center Time Measurements Across Treatment Conditions. A) Distance traveled across 5 min time periods for 60 min prior to and 90 min after saline injection, across different light cycle conditions. (Standard, n = 30, Reversed, n = 32). *: p<0.05, ** p<0.01, Two-Way ANOVA with Sidak’s. B) Total distance traveled in the same test. ***: p<0.001, Student’s t-test. C) Time in center of open field apparatus in the same test. **: p<0.01, Student’s t-test. D) Total distance traveled in an open field test from 5-155 min after drug administration to corticosterone-exposed animals, across light cycle conditions. **: p<0.01, ANOVA with Sidak’s. E) Time in center for the open field apparatus in the same test. One outlier removed from CORT + Psilocybin (ROUT). All error bars presented as Mean ± SEM.

**Figure S4.**
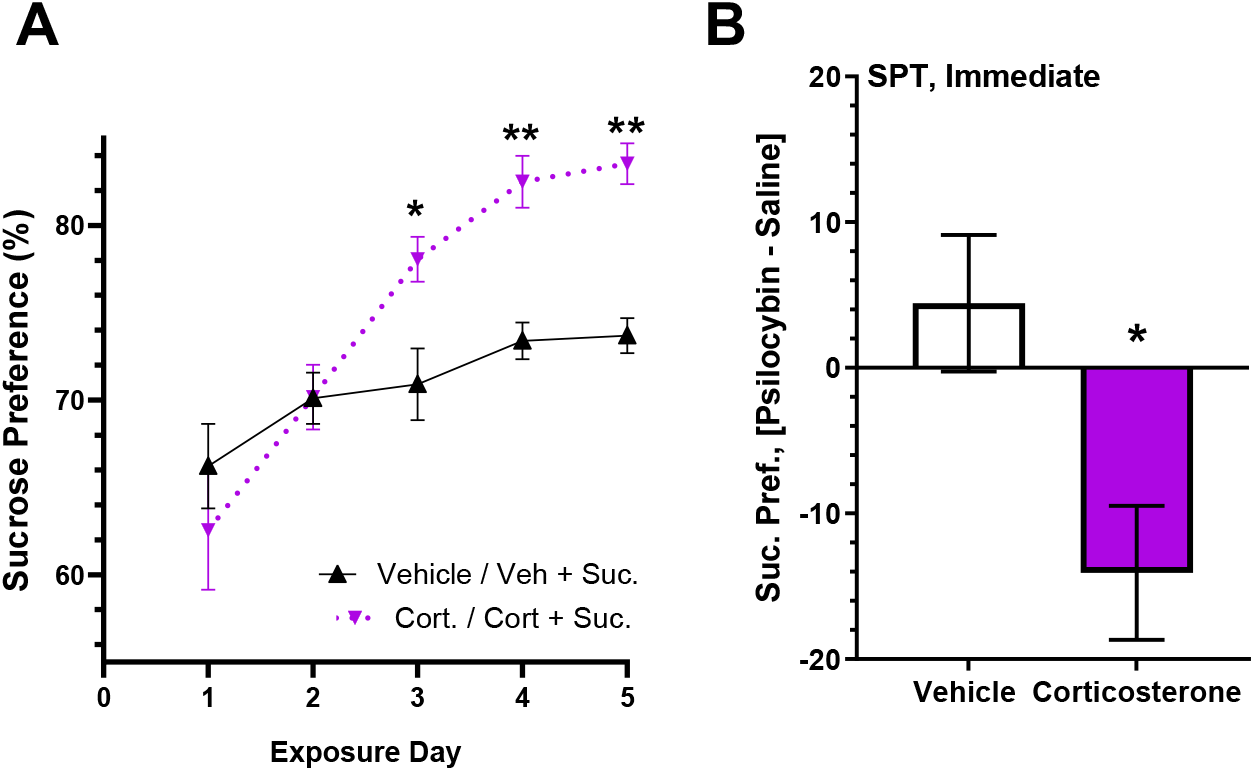
Psilocybin Acutely Blocks Avoidance of Corticosterone Water Rather Than Decreasing Preference for Sucrose Water. A) Sucrose preference on Days 1 – 5 of exposure to two-bottle choice with water containing either Vehicle / Vehicle+Sucrose or Corticosterone / Corticosterone+Sucrose (n = 15-16). * = p < 0.05, ** = p < 0.01, Two-Way RM ANOVA with Sidak’s. B) Difference in sucrose preference between animals treated with 3 mg/kg of psilocybin vs saline, starting 15 min after drug administration, during exposure to two-bottle choice with water containing either Vehicle / Vehicle+Sucrose or Corticosterone / Corticosterone+Sucrose (n = 16, vehicle; n = 8, corticosterone). * = p<0.05, Student’s t-test. All error bars presented as Mean ± SEM.

**Figure S5.**
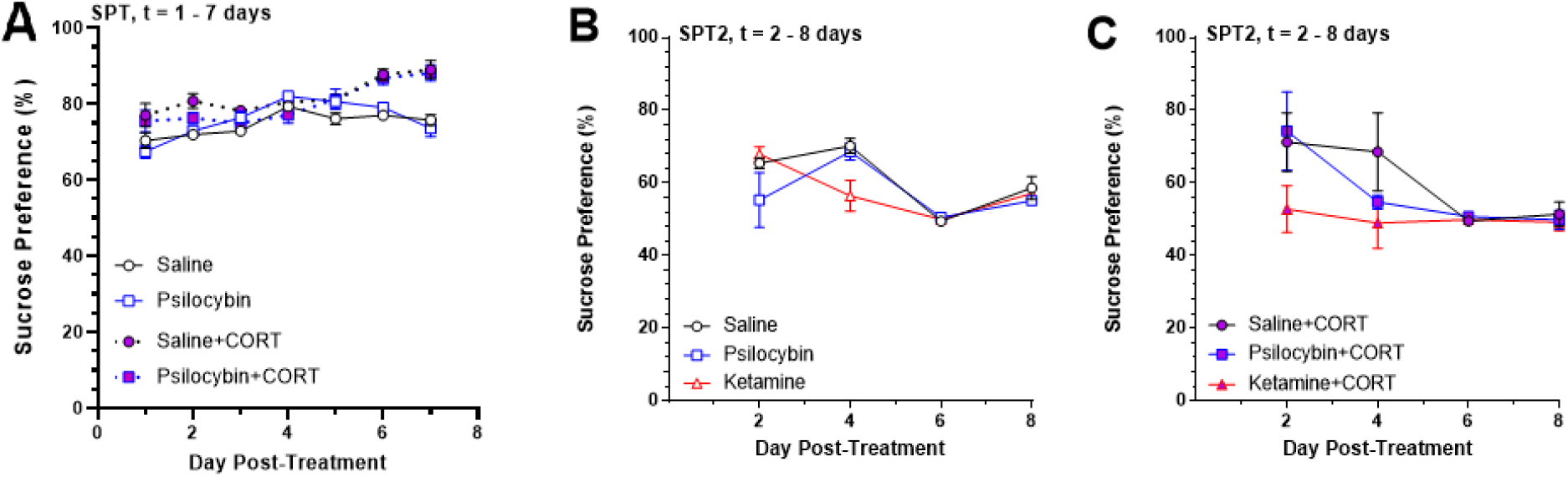
No Long-Term Effects of Psilocybin and Ketamine on Sucrose Preference. A) Sucrose preference for 1-7 days in drug-treated animals with ongoing chronic vehicle or corticosterone exposure (n=7-8). B) Sucrose preference of vehicle-treated animals upon retest beginning at 1, 3, 5, and 7 days after treatment (n= 2-3). C) Sucrose preference for 0-24 h period in drug-treated animals with prior chronic corticosterone exposure (n= 2-3). All error bars presented as Mean ± SEM.

